# Plastid double-strand RNA transgenes trigger small RNA-based gene silencing of nuclear-encoded genes

**DOI:** 10.1101/2022.12.06.519219

**Authors:** Sébastien Bélanger, Marianne C. Kramer, Hayden A. Payne, R. Keith Slotkin, Blake C. Meyers, Jeffrey M. Staub

## Abstract

Plastid transformation technology has been widely used to express traits of potential commercial importance, though the technology has been limited to traits that function while sequestered in the organelle. Prior research indicates that plastid contents can escape from the organelle, suggesting a possible mechanism for engineering plastid transgenes to function in other cellular locations. To test this hypothesis, we created tobacco plastid transformants that express a fragment of the nuclear-encoded *Phytoene desaturase* (*PDS*) gene capable of catalyzing post-transcriptional gene silencing if RNA escape to the cytoplasm occurs. We found multiple lines of direct evidence that plastid-encoded *PDS* transgenes affect nuclear *PDS* gene silencing: knockdown of the nuclear-encoded *PDS* mRNA and/or its apparent translational inhibition, biogenesis of 21-nucleotide (nt) phased small interfering RNAs (phasiRNAs), and pigment deficient plants. Furthermore, plastid-expressed double-stranded RNA (dsRNA) with no cognate nuclear-encoded pairing partner also produced abundant 21-nt phasiRNAs in the cytoplasm, demonstrating that a nuclear-encoded template is not required for siRNA biogenesis. Our results indicate that RNA escape from plastids to the cytoplasm occurs broadly, with functional consequences that include entry into the gene silencing pathway. Furthermore, we uncover a method to produce plastid-encoded traits with functions outside of the organelle and open new fields of study in plastid development, compartmentalization and small RNA biogenesis.

## INTRODUCTION

Plastid transformation technology was first developed in the model crop *Nicotiana tabacum* nearly 30 years ago (Svab et al., 1990; Svab and Maliga, 1993). Plastid transformation is an attractive technology for potential commercialization of crops engineered with biotechnology traits for several reasons: introduction of traits into the plastid genome via homologous recombination facilitates trait gene stacking, the possibility for high level transgene expression, especially in the abundant chloroplasts of leaves, and natural transgene containment due to maternal inheritance of plastids in most crop plants (Maliga, 2004; Bock, 2013; Greiner et al., 2015; Maliga and Bock, 2011). The technology has been used for expression of transgenes that impart useful traits such as insect control, herbicide tolerance, introduction of high value molecules and new metabolic pathways to enhance nutritional value (Ye et al., 2001; Dufourmantel et al., 2007; Zhang et al., 2017; Bally et al., 2018; Apel and Bock, 2009; Staub et al., 2000). Plastid transformation has been reported in numerous plant species, including commercial row crops like soybean (Dufourmantel et al., 2004), although is currently routine only in multiple Solanaceae species.

Double-stranded RNA (dsRNA) expressed in plastids has been shown to efficiently catalyze post-transcriptional gene silencing (PTGS or RNAi) in insect pests (Zhang et al., 2015; Bally et al., 2016; Jin et al., 2015; Dong et al., 2022; Wu et al., 2022a, 2022b). High-level accumulation and efficacy of long insecticidal dsRNA in plastids was attributed to the lack of dsRNA processing in plastids due to the absence of the RNAi machinery (Bally et al., 2016; Jin et al., 2015), while processing of the dsRNA to insecticidal small RNAs occurs in insect gut cells after ingestion of leaf material. The gene silencing components are encoded in the plant by nuclear genes with their activities taking place both in the nucleus and cytoplasm (Kumakura et al., 2009; Montavon et al., 2018). Typically, plant nuclear-encoded dsRNA transgenes may lose efficacy due to processing of their dsRNA in the cytoplasm to 21 – 24 nt small interfering RNAs (siRNAs), which are not efficiently taken up by insect gut cells (Li et al., 2015; Bolognesi et al., 2012). siRNA processing occurs on membrane-bound polysomes to trigger the production of secondary siRNAs that are generated on the rough endoplasmic reticulum (Li et al., 2016). These secondary siRNAs have a characteristic “phased” pattern, resulting from successive cleavage by a DICER-LIKE protein on the double-stranded RNA substrate. In many plants, DICER-LIKE4 (DCL4) generates 21-nt phased siRNAs (phasiRNAs), which are a characteristic outcome of PTGS (Liu et al., 2020).

An assumed limitation of plastid engineering is that the transgene function is sequestered in the organelle. Thus, metabolic pathways in the cytoplasm or regulatory functions in the nucleus have not been amenable to plastid engineering. Although plastids import up to ~3000 nuclear-encoded proteins (Sjuts et al., 2017; Paila et al., 2015), there is no known mechanism for plastid-encoded gene products to leave or function outside of the organelle. Despite this, there is evidence to suggest that plastid-encoded proteins and nucleic acids can be found in other cellular compartments. For example, during the natural process of chloroplast autophagy, Rubisco and other chloroplast-encoded proteins transit to or in vacuoles prior to protein degradation (Otegui, 2017; Izumi et al., 2015). Intact plastid-encoded RNAs have not been observed outside of the organelle, though plastid-encoded tRNA fragments have been observed in the cytoplasm and suggested to be involved in regulatory processes (Cognat et al., 2017; Alves and Nogueira, 2021). Over evolutionary time frames, plastid genes have migrated to the nucleus (McFadden, 2001), and can “escape” during plastid or mitochondrial transformation experiments when strong selection is used to identify rare transfer of DNA from the organelle to the nucleus (Wang et al., 2018; Fuentes et al., 2012; Stegemann and Bock, 2006; Stegemann et al., 2003; Thorsness and Fox, 1990). These observations suggest a means by which plastid DNA or RNA can exit the organelle to function in other cellular compartments.

In this study, we tested whether plastid-expressed transgenic RNAs can escape the organelle and function in the cytoplasmic post-transcriptional gene silencing pathway. We created tobacco (*Nicotiana tabacum*) plastid transformants that express a fragment of the nuclear-encoded *Phytoene desaturase* (*PDS*) gene, for which knockdown or translational inhibition of its cytoplasmic-localized mRNA results in an easily discernible, pigment deficient phenotype (Senthil□Kumar et al., 2007; Busch et al., 2002). We expressed dsRNA, sense, and antisense transcripts against *PDS* in transplastomic tobacco and found that the plastid-encoded transgenes affect nuclear *PDS* gene silencing. Highly efficient dsRNA-mediated knockdown of *PDS* was confirmed by qRT-PCR and small RNA sequencing, which indicated processing of the plastid-expressed dsRNA into 21-nt phasiRNAs, giving direct evidence for a post-transcriptional gene silencing (PTGS) mechanism. Interestingly, transplastomic lines that express dsRNA against insect gene targets, for which no pairing partner is encoded in the nuclear genome of tobacco, are also processed to phasiRNAs in the cytoplasm. Our results indicate a common process of RNA escape from plastids to the cytoplasm that can be exploited for knockdown of host nuclear-encoded genes and expands the repertoire of biotechnology tools afforded by plastid transformation technology.

## RESULTS

### Integration of a phytoene desaturase cDNA fragment in the tobacco plastid genome

We first tested the capability of a plastid-encoded fragment of the *PDS* gene to silence its nuclear-encoded, cytoplasm-localized transcript. Two genomic loci (Nitab4.5_0006338g0050.1 and Nitab4.5_0004950g0020.1; Supplemental Figure 1A), designated here as *PDS1* and *PDS2*, respectively, encoding *PDS* coding regions with >99% nucleotide sequence identity were identified in tobacco (*Nicotiana tabacum*). We selected a 294 nt region of the *PDS1* cDNA with seven nucleotide polymorphisms compared to *PDS2* to enable subsequent discrimination of the multiple *PDS* genes. Further, we introduced a small 13 nt insertion sequence into the plastid *PDS* transgenes to discriminate them from the otherwise identical nuclear-encoded *PDS* genes (Supplemental Figure 1B).

To determine if sense, antisense or dsRNA transcripts could cause silencing of the nuclear-encoded gene, we designed three plastid transgenes. The PTS40 plastid transformation vector expresses the *PDS1* cDNA fragment from two convergent plastid (P*rrn*) promoters that were previously shown to drive high-level accumulation of unprocessed dsRNA in chloroplasts (Figure 1A) (Zhang et al., 2015). As controls, both sense and antisense orientations of the *PDS1* cDNA fragment were expressed from a single plastid P*rrn* promoter and used the *E. coli rrnB* 3’-end (Zhang et al., 2015), to create PTS38 and PTS39, respectively (Figure 1A). The *PDS1* transgenes were cloned next to a selectable *aadA* spectinomycin resistance gene and between regions with identity to the tobacco plastid genome to mediate site-directed integration by homologous recombination (Figure 1A).

**Figure 1.**
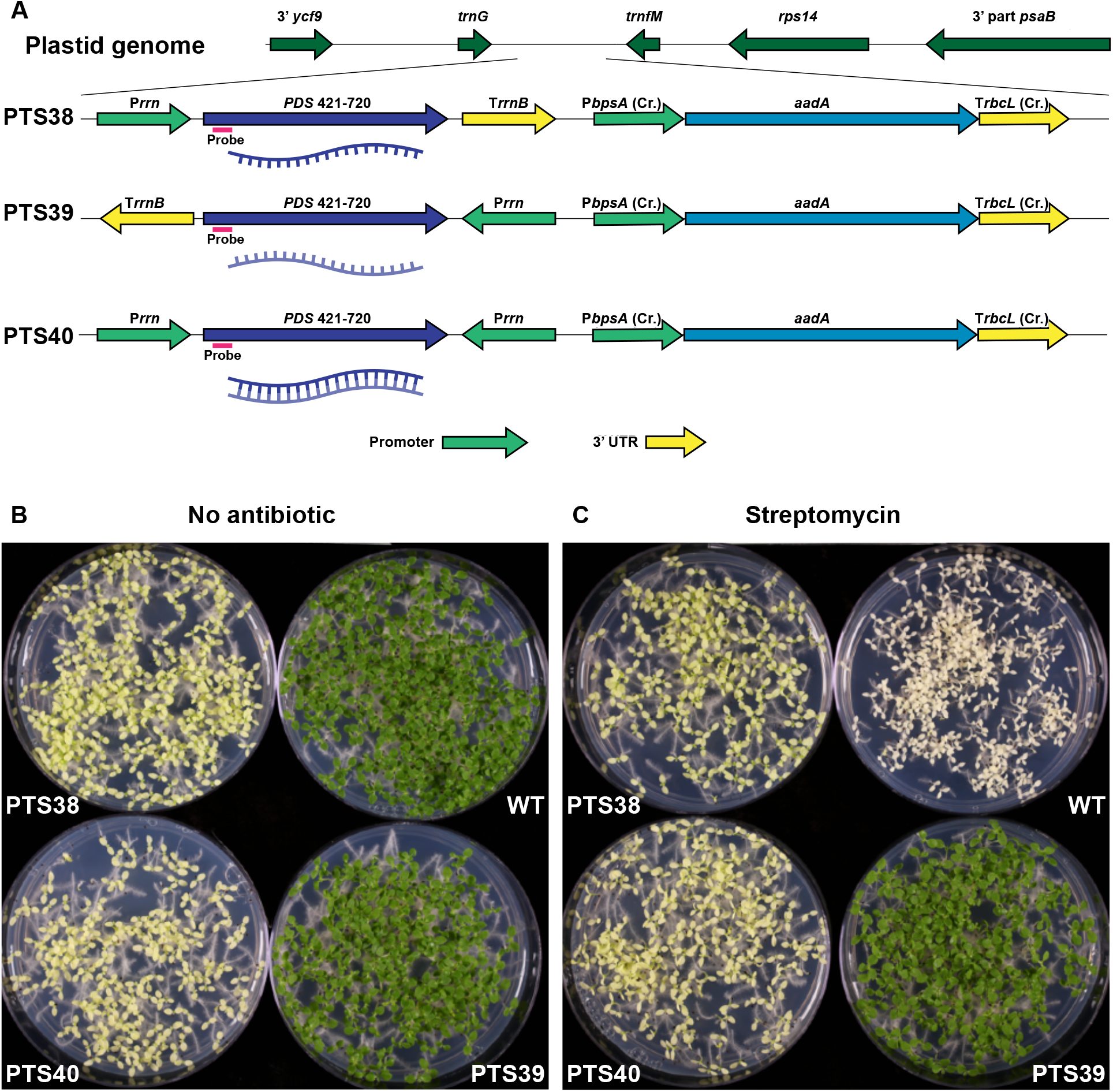
Plastid-expressed sense and double-strand RNA PDS transgenes produce pigment deficient tobacco seedlings. (A) *PDS* sense RNA (PTS38), antisense RNA (PTS39) and dsRNA (PTS40) transgenes and the *aadA* selectable marker are integrated into the plastid genome between the resident *trnG* and *trnfM* plastid genes via homologous flanking regions. Oligonucleotide probes used for *PDS* gene detection are shown by the red lines. (B, C) 12-day old T1 seedlings derived from self-fertilized T0 plants are sown on media lacking antibiotics (B) or containing spectinomycin (C). *Cr*., *Chlamydomonas reinhardtii*

Tobacco plastid transformation of *in vitro*-grown leaf tissue was used to introduce the transgenes via homologous recombination into the plastid genome. Tissue from transformed shoots was used for two subsequent rounds of plant regeneration to ensure homoplasmy of the plastid transformed lines. Integration of the transgenes was confirmed in all transformed lines during the selection process by sequencing of PCR amplicons. Site-directed integration and homoplasmy of the transgenes was confirmed by Southern blot analysis (Supplemental Figure 1C).

### Phenotype of T0 transplastomic plants

T0 plastid-transformed (transplastomic) lines grew normally in sterile tissue culture and had no apparent off-types. Homoplasmic plants were rooted in tissue culture, transferred to soil, and placed in a growth chamber with cycling light to acclimate plants for subsequent greenhouse growth. After ~10 days in soil, new leaves from PTS38 (sense) and PTS40 (dsRNA) lines, but not PTS39 (antisense) lines, were observed to be pigment deficient. The pigment-deficient phenotype appeared uniform in new leaves, though with a yellow or very pale green appearance, suggesting the presence of residual chlorophyll and/or other carotenoids (Supplemental Figure 1D). As the transplastomic plants continued to grow, subsequent new leaves were either uniformly bleached or appeared chimeric with irregular patterns of bleaching mixed with some nearly green sectors (Figure 2D). In contrast, the PTS39 *PDS* antisense line was uniformly green and had no apparent phenotype, as in wild-type controls (Supplemental Figure 1D). All independent transplastomic events for each of the constructs that were transferred to soil had similar phenotypes, indicating the observed pigment deficiency was caused by the plastid transgenes.

**Figure 2.**
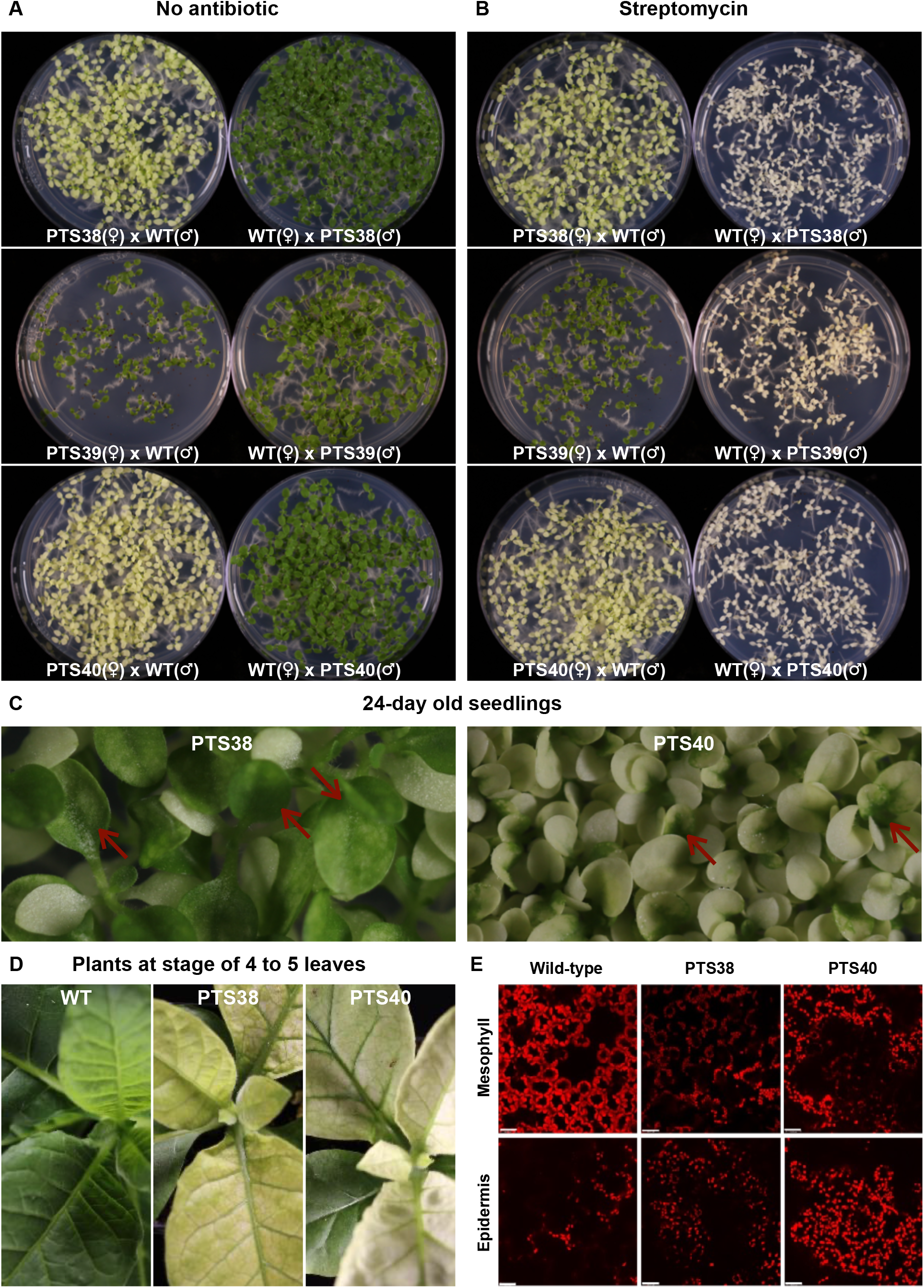
Maternal inheritance of plastid-encoded pigment deficiency. (A, B) T1 seedlings derived from reciprocal crosses of T0 transplastomic plants to wildtype plants were (e.g. WT(♀) x PTS38(♂)) sown in the absence (A) or presence (B) of spectinomycin antibiotic. (C) While cotyledons are completely bleached in the PTS38 and PTS40 seedlings, the first true leaves become green (red arrows). (D) T1 green plants placed in soil show rapid bleaching of new leaves. (E) Confocal images show size, development stage and location of plastids in mesophyll and epidermis cells in leaves of wild-type, PTS38 and PTS40 lines. The scale bar equals 20 μm. ♀, female; ♂, male.

The pigment deficiency observed in the transplastomic plants suggested a knockdown of the nuclear-encoded *PDS* gene function. Knockdown of nuclear-encoded *PDS* genes via nuclear transformation technologies typically results in albino leaf tissues (for example, Senthil□Kumar et al., 2007) due to lack of carotenoid accumulation and concomitant degradation of chlorophyll, suggesting some differences between plastid-expressed *PDS* gene silencing and the previously observed nuclear gene silencing approaches. Although the pigment-deficient lines grew much more slowly than green plants, they eventually out grew the bleaching phenotype and were transferred to the greenhouse, where lateral branching occurred, and flowering was ultimately profuse with an apparently normal seed set.

### Inheritance of plastid-encoded traits

To confirm maternal inheritance of the plastid-encoded traits, antibiotic resistance and pigment deficient phenotypes of seedlings from self-fertilized transplastomic lines were analyzed. As expected, wild-type seedlings were green on medium lacking antibiotics (Figure 1B) but were uniformly bleached white on medium containing spectinomycin (Figure 1C), indicating their sensitivity to the antibiotic. In contrast, seedlings derived from self-fertilization of the PTS38 and PTS40 lines had an intermediate phenotype, uniformly bleached yellow rather than white when grown on plates with or without spectinomycin (Figure 1B–C). These results indicate that the PTS38 and PTS40 lines are resistant to the antibiotic and homoplasmic, and that the pigment deficiency may be due to silencing of the nuclear-encoded *PDS* genes. Seedlings of the PTS39 antisense line were uniformly green, indicating uniform resistance to the antibiotic, as expected for a homoplasmic, plastid-encoded trait, but no pigment deficiency was observed in these lines.

Reciprocal crosses of plastid-transformed lines with wild-type plants are used to prove maternal inheritance of a plastid transgenic trait (Svab et al., 1990; Staub and Maliga, 1992). When the PTS38 and PTS40 plants were used as female parents in crosses to wild-type plants, all seedlings were uniformly pigment deficient on media lacking antibiotics (Figure 2A). In contrast, when the plastid-transformed lines were used as the male pollen donor in crosses to wild-type plants, all seedlings were uniformly green (Figure 2A). In the presence of the selective antibiotic (Figure 2B), all female-derived transplastomic seedlings were again pigment deficient but uniformly antibiotic resistant, while all seedlings were uniformly sensitive and bleached white when the plastid-transformed lines were used as the pollen donor to wild-type plants. These results confirmed that the pigment deficient phenotype and antibiotic resistance is maternally inherited and not transmitted through pollen, as expected for plastid-encoded traits. Maternal inheritance was further confirmed in T2 and T3 generations of the PTS40 and PTS38 lines, confirming that there is no active copy of the *PDS* transgenes in the nuclear genome.

Although PTS38 and PTS40 T1 seedling cotyledons were uniformly pigment deficient, the first true leaves emerged as partly green and subsequent leaves were uniformly green. However, the pigment deficient phenotype of PTS40 seedlings is stronger and persists longer than in the PTS38 line, as evidenced by more rapid greening of its first true leaves (Figure 2C). Interestingly, when PTS38 and PTS40 T1 plants were grown in tissue culture to the 4 to 5 leaf stage and green plants were then transferred back to soil, the chlorophyll-deficient phenotype of newly emerged leaves quickly returned (Figure 2D). These results indicate that the pigment deficiency is subject to developmental timing.

### Chloroplast division is blocked in a tissue-specific manner in chlorophyll deficient lines

To gain insight into possible defects in plastid development that could help potentiate *PDS* gene silencing, plastid morphology in leaves from T1 plants grown in soil was examined by confocal microscopy. Mesophyll cells of wild-type plants (and PTS39 lines, data not shown) have characteristic large chloroplasts (average size ~5 μM) arranged along the periphery of each cell (Figure 2E). Chloroplasts appeared fully developed and densely packed, side-by-side. In contrast, PTS38 and PTS40 transplastomic lines contain smaller (average size ~2.5 μM) plastids, more loosely arranged, mostly along the periphery of the cell. Interestingly, most of the plastids appeared as closely associated pairs, suggesting a block in plastid development shortly after plastid division. In contrast to chloroplasts of mesophyll cells, epidermal cell plastids were observed to be relatively small (~3 μM average size) and loosely arranged along the cell periphery in wild-type, PTS38 and PTS40 lines. These results suggested that the *PDS* gene knockdown may not occur in epidermal cells of the plastid transformed lines, allowing some chlorophyll to accumulate in those cells, resulting in the yellow appearance of PTS38 and PTS40 lines. Moreover, although a block in plastid division is apparent, it’s unclear if this can contribute to leakage of plastid transcripts into the cytoplasm.

### qRT-PCR confirms knockdown of PDS mRNA in plastid transformed lines

To confirm that pigment deficiency of plastid transformed lines is due to silencing of the nuclear-encoded, cytoplasm-localized *PDS1* and/or *PDS2* mRNAs, quantitative reverse-transcriptase PCR (qRT-PCR)-mediated analysis of total cellular RNA from T1 transplastomic and wild-type seedlings was performed. PCR primer pairs that map outside of the plastid-encoded *PDS* transgene (Supplemental Figure 1) were used to distinguish and quantify only the endogenous nuclear-encoded *PDS* mRNAs. In 12-day-old seedlings, when cotyledons of PTS38 and PTS40 lines are uniformly bleached, the level of *PDS1* mRNA is significantly reduced to about half and *PDS2* mRNA is reduced slightly in the PTS40 dsRNA lines compared wild-type. In contrast, *PDS1 and PDS2* mRNA is not reduced in the PTS38 and PTS39 plastid-transformed lines (Figure 3A). At the later developmental stage (24-day-old seedlings) when the first true leaves of PTS40 seedlings have begun to turn green (Figure 2C), nuclear-encoded *PDS1* mRNA levels recover to near wild-type levels, consistent with the cessation of the pigment-deficient phenotype (Figure 3B). These results confirm that the pigment-deficient phenotype of PTS40 lines is due to knockdown of the nuclear-encoded *PDS* mRNA, while the bleaching in the PTS38 sense lines is apparently due to a different gene silencing mechanism.

**Figure 3:**
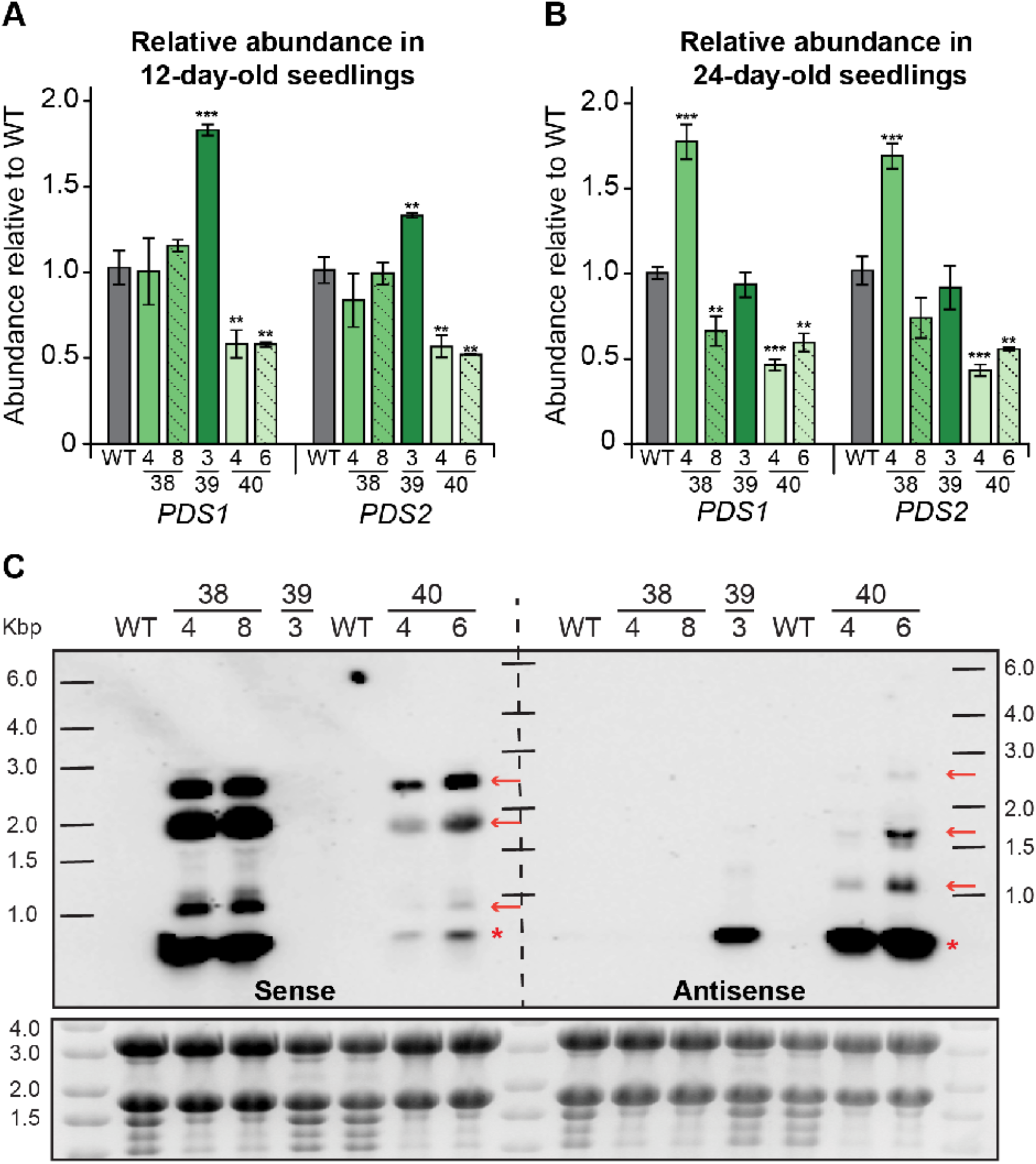
Abundance and expression pattern of nuclear- and plastid-encoded PDS genes. (A-B) Relative abundance of nuclear-encoded *PDS1* and *PDS2*, as measured by qRT-PCR, in 12-day-old (A) and 24-day-old (B) seedlings from wild-type and PTS38, PTS39 and PTS40 transplastomic lines. All RNA expression values are relative to wild-type and normalized to the housekeeping gene, *Actin*. Error bars represent standard error around the mean. *, **, *** denotes p-value < 0.05, 0.01, 0.001, respectively, Student’s t-test. (C) Northern blot with wild-type and independent transplastomic events of PTS38, PTS39 and PTS40 lines. Membrane was either blotted with a probe recognizing the sense strand of *PDS1* (left) or antisense strand of *PDS1* (right). Ladder is RiboRuler High Range RNA ladder. Gel image before transfer is represented below the blot. Note that transcripts for the endogenous nuclear-encoded *PDS1* and *PDS2* genes are expected to be expressed at much lower levels than the plastid transgenes and are not observed on the northern blot.

### Northern blot analysis confirms high-level expression of plastid-encoded PDS transcripts

Previous northern blot studies showed high-level dsRNA accumulation in plastids, though the relative accumulation of sense and antisense strands expressed from convergent promoters was not examined (Zhang et al., 2015). Using oligonucleotide probes from within the *PDS1* plastid transgene fragment (Figure 1), accumulation of plastid-encoded *PDS1* sense or antisense (Figure 3C) RNA from the transplastomic lines was examined. The PTS38 line accumulates very high levels of sense strand transcripts of the expected ~294 nt *PDS1* fragment size and several higher molecular weight transcripts, which are likely due to inefficient transcriptional termination, as is often observed from plastid genes (Bock, 2013). The PTS40 line accumulates similar sized sense-strand transcripts as the PTS38 line, albeit at much lower levels. Interestingly, the PTS40 line accumulates higher amounts of the *PDS1* ~294 nt fragment antisense transcript, suggesting that unpaired sense and antisense transcripts likely exist in this line, in addition to dsRNA. Readthrough transcripts also accumulate to low levels in this line, suggesting a difference in mRNA stability between sense and antisense transcripts in the PTS40 line. In contrast to the other transplastomic lines, the PTS39 line accumulates only the ~294 nt sense-strand *PDS1* fragment transcript and no apparent readthrough transcripts.

### Processing of plastid transgene-encoded small in the cytoplasm

Reduction of nuclear-encoded *PDS* mRNA levels via plastid transgenic RNAs suggests that the latter can enter the PTGS pathway, which is characterized by the presence of 21 to 24 nt siRNAs. Since previous reports of chloroplast-transformed plants carrying dsRNA transgenes did not observe siRNA accumulation by northern blot analysis (Zhang et al., 2015; Bally et al., 2016), we used the higher-resolution technique of small RNA deep sequencing of the 12-day old transplastomic and wild-type seedlings derived from self-pollinated plants. As expected, no siRNAs mapping to either *PDS1* or *PDS2* nuclear genes were detected in wild-type seedlings (Table 1). In contrast, abundant siRNAs (~180 to 280 reads per million, RPM) that match *PDS* were observed in all PTS38 and PTS40 (bleached) lines (Figure 4 and Table 1). Interestingly, one (green) PTS39 line accumulated detectable siRNA reads, though to a much lower degree (~20 RPM; Supplementary Figure 2). Inspection of the siRNA sequences indicates that none mapped to polymorphic *PDS2* sequences (Supplementary Figure 3), and no secondary siRNAs derived from the nuclear-encoded *PDS* genes beyond the 294 nt plastid *PDS* transgene region were observed, suggesting that the siRNAs derive directly from plastid transcripts.

**Figure 4.**
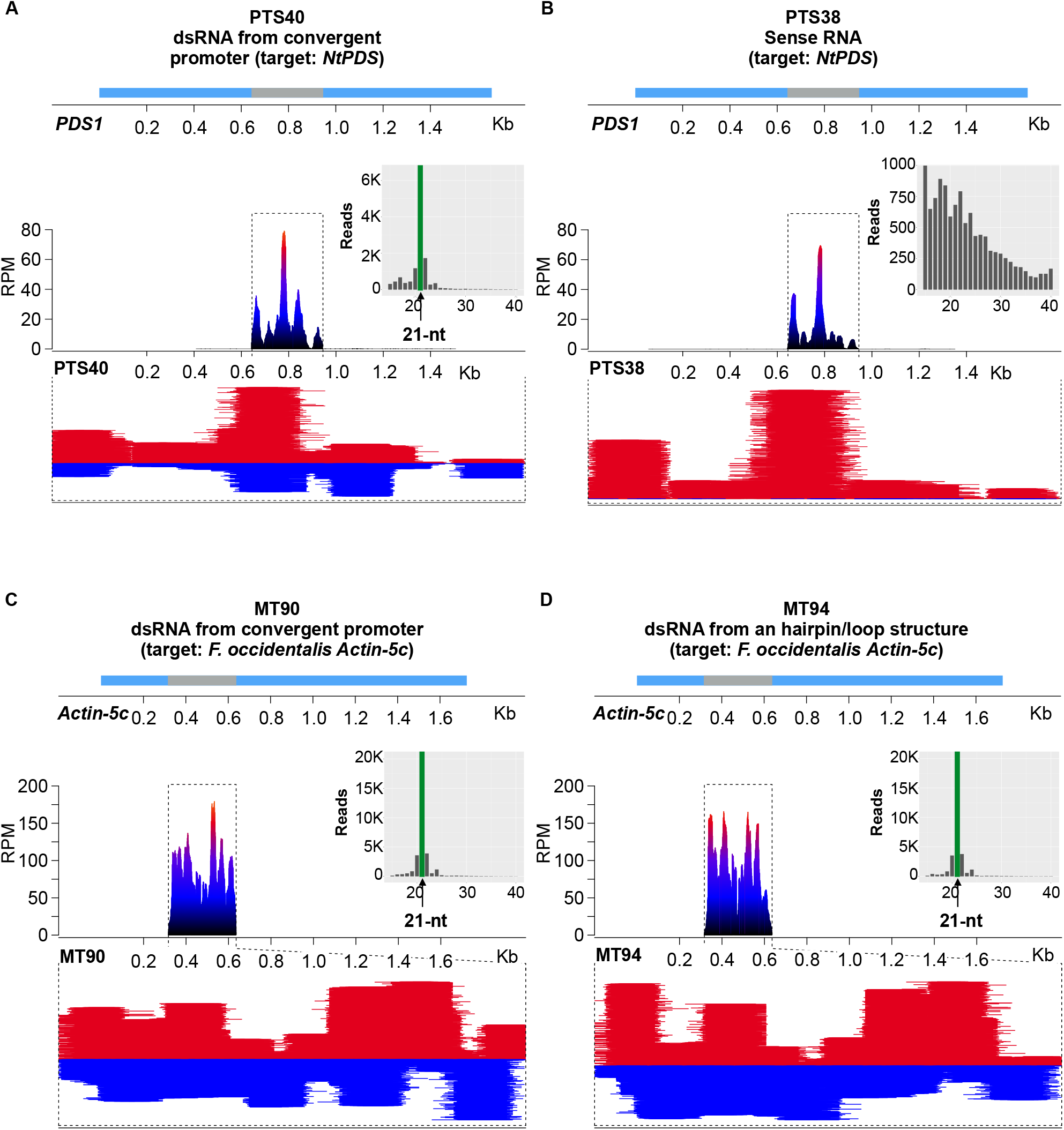
Plastid-expressed dsRNA is processed to 21-nt phasiRNAs. (A-B) Mapping and accumulation of *PDS1* siRNAs (red, sense strand; blue, antisense strand) in the PTS38 and PTS40 transplastomic lines. (A) Accumulation of siRNAs in the PTS38 line map to the PDS1 sense strand and show no bias in length (insert) whereas (B) siRNAs of predominantly 21-nt (insert) accumulate on both strands of *PDS1* in the PTS40 lines. The relative location of the *PDS* transgene fragment (grey) in the nuclear-encoded *PDS1* gene (light blue) are represented above the panels. (C-D) Mapping and accumulation of siRNAs in the MT90 and MT94 lines expressing the *Frankliniella occidentalis Actin-5c* gene from either convergent promoters (C) or a hairpin/loop RNA (D) transgene. Inserts show the length distribution of reads showing predominantly 21 nt siRNAs. The relative location of the transgene fragment (grey) in the *Actin-5C* gene (light blue) are represented above the panels. The y-axis indicates the distribution of read abundance normalized in reads per million (RPM).

**Table 1.** Plant lines, phenotype and siRNA characterization.

| Plant line | Expression strategy | Target gene | Phenotype | RPM | Major RNA length | Phase score |
| --- | --- | --- | --- | --- | --- | --- |
| PTS38-4 | Sense RNA | PDS | Bleached | 198 | 15 | - |
| PTS38-8 | Sense RNA | PDS | Bleached | 179.8 | 19 | - |
| PTS39-3 | Antisense RNA | PDS | Green | 19.6 | 21 | 17.3 |
| PTS39-4 | Antisense RNA | PDS | Green | - | - | - |
| PTS40-4 | Convergent promoter | PDS | Bleached | 280.5 | 21 | 35 |
| PTS40-6 | Convergent promoter | PDS | Bleached | 260.2 | 21 | 35.3 |
| MT90-2 | Convergent promoter | Actin-5c | Green | 691.7 | 21 | 39.7 |
| MT90-5 | Convergent promoter | Actin-5c | Green | 1,295.40 | 21 | 61.7 |
| MT91-2 | Convergent promoter | SNF7 | Green | 71.1 | 21 | 6.5 |
| MT91-4 | Convergent promoter | SNF7 | Green | 106.8 | 21 | 10.2 |
| MT94-1 | Hairpin/loop RNA | Actin-5c | Green | 832.4 | 21 | 57.7 |
| MT94-2 | Hairpin/loop RNA | Actin-5c | Green | 1,331.60 | 21 | 51 |
| MT95-1 | Hairpin/loop RNA | SNF7 | Green | 186.4 | 21 | 52.8 |
| MT95-2 | Hairpin/loop RNA | SNF7 | Green | 145.6 | 21 | 27.4 |
| WT | N/A | N/A | Green | - | - | - |

Note: A dash indicates either no reads were detected, or no phasing score was observed.

To provide further insights into the mechanism of siRNA biogenesis from the plastid transgenes, we examined their size distribution, strandedness, and potential “phasing”. siRNA phasing is an indication of recursive Dicer processing and may be amplified via an RNA-dependent RNA polymerase (RDR protein) acting in the cytoplasm (Liu et al., 2020). For the PTS40 dsRNA lines, siRNAs map abundantly to both strands of the *PDS1* transgene and 21-nt siRNAs make up the vast majority of reads across both strands (Figure 4A and insert). The phasing analysis for dsRNA PTS40 lines indicated a highly significant phasing score of ~35 (Table 1). These results indicate that Dicer-like processing resulting in phasiRNAs occurs from plastid-transgene RNAs, and such processing should localize in the cytoplasm of transplastomic plants. Abundant siRNAs were also mapped to *PDS1* in the transplastomic PTS38 sense strand lines (Figure 4B). However, in contrast to the PTS40 dsRNA lines, PTS38-derived siRNAs mapped exclusively to the sense strand of the *PDS1* gene, have a broad size range distribution with no discernible size peak and were not phased. These latter results suggest PTS38-derived siRNAs may result from transcript degradation in the cytoplasm or the chloroplast, rather than Dicer-dependent cleavage. Taken together with the qRT-PCR results that indicates no change of nuclear-encoded PDS mRNAs (Figure 3) and no apparent phasiRNAs, these data suggest that the bleaching phenotype observed in the PTS38 line is likely due to a different mechanism than in the PTS40 lines, such as translational inhibition of PDS.

Accumulation of 21-nt phasiRNAs in the PTS40 dsRNA line presumably results from processing of plastid-derived transcripts in the cytoplasm after RNA movement by unknown mechanisms. To rule out previously uncharacterized RNA processing events of transgenic transcripts inside the plastid, we prepared small RNA libraries from purified plastids from the same 12-day old transplastomic T1 seedlings as described above. In all transplastomic lines, plastid-localized small RNAs were abundant with a broad size distribution, but no clear peak at 21-nt or phasiRNAs were observed (Supplemental Figure 4). These results confirm that processing of plastid transcripts to phasiRNAs occurred in the cytoplasm and not in the organelle.

### Processing of plastid-transgene-derived dsRNA to 21-nt phasiRNAs is a universal phenomenon

The finding of abundant 21-nt phasiRNAs derived from the plastid *PDS1* raised the question of whether a nuclear-encoded complementary target RNA is required for processing into siRNAs. To examine this question, we utilized transplastomic tobacco lines that express dsRNA against non-plant (insect) targets, from either two convergent plastid promoters identical to the *PDS* dsRNA transgene (MT90 and MT91) or via a single chloroplast promoter driving a hairpin/loop dsRNA construct (MT94 and MT95) (Figure 4 and Supplementary Figure 5). The constructs were designed to express dsRNA against two insect gene targets, *SNF7* and *Actin5C* from *Frankliniella occidentalis* (Wu et al., 2022a).

We again performed small RNA sequencing of total cellular RNA from 12-day old transplastomic seedlings The results were similar for the convergent chloroplast promoters and the hairpin/loop dsRNA constructs, and across two independent transplastomic lines tested for each construct (Figure 4C and 4D; Table 1). In all cases, siRNAs were abundant and mapped across both DNA strands of *SNF7* and *Actin5C*, and along the entire genic region that comprises the plastid transgenes, with no apparent transitive siRNAs mapping outside of this region. Importantly, the size distribution of the siRNAs showed a strong peak at 21-nt in all events and each event had a highly significant phasing score from ~27 – 61 (Table 1). These results confirmed that plastid-expressed dsRNA enters the host cytoplasmic RNAi pathway to generate 21 nt phasiRNAs, are derived from the transgene, and arise independent of complementary nuclear-encoded sequences.

## DISCUSSION

The ability of plastid-encoded RNAs to enter the gene silencing pathway in the cytoplasm is a surprising finding. We observed that plastid-encoded PDS RNAs affect nuclear PDS gene silencing via multiple lines of evidence, including accumulation of 21-nt phasiRNAs that derive from Dicer-like processing of dsRNA in the PTS40 and MT lines (Figure 4). Interestingly, lack of siRNAs spreading to adjacent PDS1 sequences or siRNAs derived from the PDS2 polymorphic regions suggests that plastid-encoded transcripts can be used directly in the cytoplasm as template for Dicer-like 4 cleavage without RDR activity. In support of this conclusion, the processing of plastid transgene-encoded dsRNAs to 21-nt phasiRNAs does not require a plant host-encoded transcript target to enter the RNAi pathway, as evidenced by abundant phasiRNAs derived from the dsRNA plastid transgenes encoding an insect gene fragment with no cognate nuclear-encoded complimentary partner. Our data thus suggest a departure from the current understanding of plant phasiRNA biogenesis, which typically initiates with an Argonaute-catalyzed (AGO) cleavage of a single-stranded mRNA precursor, then converted to dsRNA by an RDR protein, followed by processing into 21-nt or 24-nt RNA duplexes by a Dicer-like (DCL) protein (Hung and Slotkin, 2021; Liu et al., 2020; Fei et al., 2013). On the other hand, we also observed apparent translational inhibition from sense-strand RNAs in the PTS38 line, as evidenced by pigment deficient plants with no PDS RNA knockdown via qRT-PCR (Figure 3) or phasiRNAs by small RNA sequencing (Figure 4). Translational repression of nuclear genes may occur by interfering with translationally active ribosomes via multiple mechanisms in response to accumulation of translatable transcripts at high concentrations (Li et al., 2013) as is observed in the PTS38 line by northern blot (Figure 2). However, future work will be needed to determine the mechanism acting from plastid-expressed sense RNAs.

Movement of the plastid transcripts to the cytoplasm is a prerequisite to phasiRNA production and nuclear-encoded *PDS* gene silencing since the PTGS machinery does not exist in plastids (Zhang et al., 2015; Bally et al., 2016). Our results indicate that movement of transcripts out of the plastid compartment must happen commonly, though may be developmentally regulated, as evidenced by the changing nature of the pigment deficient phenotype. The initial triggering events originate in germinating seedlings and last until the first true leaves emerge partly green, and subsequent plant growth in tissue culture looks normal. Completely green and healthy plants in tissue culture again show bleaching of new leaves when plants are transferred to soil. Bleaching of plant leaves in soil persists for several weeks, suggesting a continual escape of plastid-expressed *PDS* transcripts and continual entry into the gene silencing machinery in the cytoplasm. These results suggest a continual turnover of a subpopulation of plastids during plant development, reminiscent of autophagic turnover of plastids (reviewed in (Izumi et al., 2019; Woodson, 2019; Izumi and Nakamura, 2018; Zhuang and Jiang, 2019)), perhaps initially in response to limited photosynthetic activity that induces sugar starvation. We speculate that plastid redox or other signals accumulating during these developmental stages may trigger plastid autophagy, liberating their contents to the cytoplasm *en route* to the vacuole by unknown means. During this process, highly expressed plastid RNAs are apparently stable enough to be captured by the PTGS apparatus located in the cytoplasm.

Liberation of plastid-derived transcripts to the cytoplasm occurred in the absence of any specific treatments designed to facilitate the process. Our observations are different from previous reports of “escape” of DNA from mitochondria or chloroplasts in which a strong selection is required to identify rare transfer of a chloroplast transgene to the nucleus (Thorsness and Weber, 1996; Bock, 2015; Huang et al., 2003; Bock and Timmis, 2008). Likewise, our work contrasts with a report that utilized paraquat herbicide or bacterial infection to catalyze reactive oxygen species that disrupt chloroplast membranes, thus allowing leakage of a chloroplast-localized fluorescent protein to the cytoplasm (Kwon et al., 2013). It will be interesting to determine if a chloroplast-expressed transgenic protein can also move out of the chloroplast to catalyze effects in other cellular compartments or provide for useful traits thought previously not to be amenable to chloroplast transformation technology.

While the precise molecular mechanisms leading to small RNA production from plastid transgenes and activity in the cytoplasm against a nuclear-encoded gene will require additional research to fully understand, our results have implications for expanding the range of targets available for plastid engineering. Recent studies indicate that transplastomic plants accumulate high levels of long unprocessed dsRNA and are effective against multiple insect pests. In each of these reports, no apparent small RNA production from chloroplast transgenes was observed via northern blots, leading the authors to conclude that DCL proteins and the rest of the RNAi machinery are absent from plastids (Zhang et al., 2015; Bally et al., 2016). Our data suggest that the lack of processing of long dsRNA to siRNA in plastids may not be a limitation for controlling plant pathogens that preferentially take up siRNAs or don’t digest plant tissues, for example, viral, nematode and some fungal and oomycete pathogens (Liu et al., 2019; Jaubert-Possamai et al., 2019; Qiao et al., 2021; Taliansky et al., 2021). The ability to now silence plant nuclear genes from the plastid enables engineering of an array of traits previously unavailable to plastid transformation technology, including metabolic processes, biomass, and grain yield (Mansoor et al., 2006; Kamthan et al., 2015; Saurabh et al., 2014). The results reported here also add a new potential pathway for chloroplast-to-nucleus signaling that has not been previously considered (reviewed in Jan et al., 2022; Liebers et al., 2022).

## ACKNOWLEDGMENTS

We thank Dr. Ralph Bock for the gift of plasmid pJZ199 and for seeds of tobacco MT lines carrying the insect control genes. We also thank and acknowledge Dr. Kirk Czymmek for imaging support from the Advanced Bioimaging Laboratory (RRID:SCR_018951) at the Donald Danforth Plant Science Center and usage of the Leica SP8-X confocal microscope acquired through an NSF Major Research Instrumentation grant (DBI-1337680). J.M.S. thanks members of Plastomics, Inc for providing tissue culture media and other technical assistance during various stages of this work.

## AUTHOR CONTRIBUTIONS

J.M.S. conceived of the project, performed molecular cloning, generation and phenotyping of transgenic PDS tobacco lines. B.C.M., R.K.S, J.M.S., S.B. and M.C.K. designed the RNA studies; qRT-PCR and northern blotting was performed by M.C.K. and H.A.P.; small RNA and data analysis: S.B., J.M.S., B.C.M, R.K.S and M.C.K.; S.B. and J.M.S. wrote the manuscript with contributions and editing by all authors.

## MATERIALS AND METHODS

### Construction of plastid transformation vectors

The *PDS1* gene fragment was synthesized and cloned as a KpnI/SbfI DNA fragment between the convergent P*rrn* promoters in vector pJZ199 (Zhang et al., 2015). In pPTS38, the *E. coli rrnB* terminator fragment was synthesized and cloned as a KpnI/NotI fragment to replace one of the P*rrn* promoters, to create the sense *PDS* construct. In pPTS39, the *rrnB* terminator was cloned as an SbfI/SalI fragment to replace the opposite P*rrn* promoter, creating the antisense *PDS* construct. The *PDS* transgenes were cloned next to a chimeric *aadA* spectinomycin resistance gene driven by *Chlamydomonas reinhardtii* chloroplast *psbA* gene promoter and *rbcL* gene 3’-untranslated region. The *PDS* and *aadA* transgenes are flanked by regions of identity (~1950 nt and ~670 nts) to the tobacco chloroplast genome, resulting in integration of both transgenes between the resident *trnfM* and *trnG* chloroplast genes in plastid transformed plants.

### Plant growth, transformation, and selection of chloroplast transformants

*Nicotiana tabacum* cv Petit Havana plants were grown aseptically from seedlings on MS agar medium for ~4 weeks at 28 C in 16 hr. light/8 hr. dark cycling light (using 4500K cool fluorescent bulbs). Young leaves were harvested for particle bombardment, placed abaxial side up and bombarded using the BioRad PDS1000 He gun according to standard procedures (Maliga and Svab, 2010). Transplastomic events were selected by growth on 500 mg/L spectinomycin. Primary transformants typically arise as shoots on this medium; young leaf tissue from shoots is dissected and used for a second and subsequently repeated for a third round of plant regeneration on selective medium to ensure homoplasmy of the plastid transformed lines. Plastid transformants are confirmed by PCR-sequencing of amplification products to confirm transgene insertion and identity in each transformed line.

Plastid transformed lines were rooted on MS medium containing 500 mg/L spectinomycin and allowed to grow to the 4-5 leaf stage before transfer to soil. For reciprocal crosses of plastid transformed lines to wild-type plants, flowers were emasculated by hand and manually pollinated, then individually bagged until seed pods are mature.

Seeds from wild-type and plastid-transformed plants were surface sterilized using 10% Clorox solution with a few drops of Tween-20 for 10 minutes with shaking. After sterilization, seeds were washed with at least four changes of excess sterile water and sown on agar medium at 24 °C with 16 hours light for 12 days prior to harvest. Two events per construct were grown for isolation of whole-cell and chloroplast-enriched RNA fractions.

### Confocal microscopy

For imaging of leaf tissues, samples were syringe infiltrated with water before microscopic analysis and imaged on a Leica SP8-X (Leica Microsystems, Wetzlar Germany) with a water immersion 63x HC Plan Apochromat CS2 objective lens (numerical aperture 1.2). Cell wall autofluorescence was visualized using 405 nm diode laser excitation, 414-516 nm emission and a HyD detector. Chlorophyll autofluorescence was visualized using 649 nm laser excitation, 658-768 nm emission and photomultiplier tube detector (PMT). Three-dimensional (3D) image stacks were acquired with a pinhole of 1 airy unit, 512×512 pixel images, field-of-view of 184.52 μm^2^ and 0.9 μm z-interval. Z-stacks were rendered as a 3D maximum intensity projection. Confocal z-stack were imported into ImageJ (version 1.53c) and the line profile tool was used to measure the long axis of individual chloroplasts from the epidermis, mesophyll and stomata. Line measurements were exported into an excel spreadsheet using the Analyze and Measure feature of ImageJ, and box plots and median plastid size was calculated from at least 50 measurements in each image.

### qRT-PCR of PDS1 and PDS2 levels in transgenic plants

To examine levels of *PDS1* and *PDS2* in PTS38, 39, and 40, seedlings were collected 12 days and 4 weeks after sowing and directly frozen in liquid nitrogen. Tissue was then pulverized in liquid nitrogen and transferred to TRIzol Reagent (Invitrogen, Carlsbad, CA) and RNA was isolated according to the manufacturer’s instructions. RNA was then treated with RNase-free DNase (Qiagen, Valencia, CA) for 25 minutes at room temperature, ethanol precipitated and resuspended in nuclease-free water. Reverse transcription (RT) was performed using Advantage RT-for-PCR kit (Takara, San Jose, CA) following the manufacturer’s instructions with 20 μM oligo (dT)_18_. cDNA was diluted 1:10 in nuclease-free water for qRT-PCR.

qRT-PCR was performed using 2X PowerUp SYBR Green Master Mix (Applied Biosystems, Carlsbad, CA) as follows per well: 5 μL 2X PowerUp SYBR Green Master Mix, 0.75 μL cDNA (diluted 1:10), 1.25 μL nuclease-free water, and 3 μL combined 1.5 μM forward and reverse primers. All qRT-PCR reactions were performed in three technical replicates and all primers were tested for non-specific amplification using water and specificity using genomic DNA and analysis of melt curves. All reactions were run using the following program: 95°C for 10 minutes; 40 cycles of 95°C 30 sec, 55°C 30 sec, 72°C 30 sec. Melt curves were generated by heating the final PCR 1.6°C/s to 95°C for 15 sec, decreasing the temperature to 60°C at 1.6°C/s and slowly increasing back to 95°C at 0.1°C/s. All primers are listed in Supplemental Table 1. Results were first normalized to the housekeeping gene *Actin* (dC_T_) and then normalized to wild-type (ddC_T_). Relative quantity of ddC_T_ value was calculated as 2^-(ddC_T_)^. Error bars represent standard error of the mean (SEM).

### Northern blot analysis

For the denaturing agarose northern blot, 20 μg of DNase-treated total RNA was diluted to a final volume of 24.5 μL in nuclease-free water. 2X NorthernMax Formaldehyde Load Dye (Invitrogen, Carlsbad, CA) was added to the RNA samples and 10 μL RiboRuler High Range RNA Ladder (ThermoScientific, Carlsbad, CA) and denatured for 15 minutes at 65°C. RNA was immediately placed on ice and 1μL ethidium bromide (20 mg/mL) was added and mixed well. Probes were prepared using the Biotin 3’ End DNA Labeling Kit (ThermoScientific, Rockford, IL) as described by the manufacturer’s instructions. Probes were synthesized by IDT (Newark, NJ) and are listed in Supplemental Table 1.

To make the denaturing agarose gel, UltraPure agarose (Invitrogen, Carlsbad, CA) was dissolved in water, allowed to cool to ~60°C and 10X NorthernMax Denaturing Gel Buffer (Invitrogen, Carlsbad, CA) (pre-warmed to 60°C) was added to have 1.2% agarose gel. 10 μg of RNA was loaded into each well (each sample was split between two wells) and run in 1X MOPS Electrophoresis Buffer (Fisher Bioreagents) for 2.5 hours at 95V. The gel was imaged and washed 2 x 15 minutes in nuclease-free water then 2 x 15 minutes in 10X UltraPure SSC Buffer (Invitrogen, Carlsbad, CA) with gentle shaking. Samples were transferred to BrightStart - Plus membrane (Invitrogen, Carlsbad, CA) membrane using capillary action in 10X UltraPure SSC Buffer overnight. The next day, RNA was crosslinked to the membrane using Stratagene UV Stratalinker 2400 and incubated for 30 minutes in pre-warmed NorthernMax Prehyb/Hyb Buffer (Invitrogen, Carlsbad, CA) at 42°C in a hybridization oven with rotation. 20 μL of biotinylated probes were added to Prehyb/Hyb solution and incubated overnight at 42°C with rotation. The next day, the probe was removed and the membranes were washed in 2X SSC, 0.5% SDS wash solution 2 x 30 minutes. The membranes were then treated with the Chemiluminescent Nucleic Acid Detection Module (ThermoScientific, Carlsbad, CA) for detection, according to the manufacturer’s instructions. Membranes were visualized using BioRad ChemiDoc XRS+.

### Southern blot analysis of plastid transformed lines

Leaves from plants or seedlings grown aseptically in tissue culture were used for total cellular DNA isolation using the DNAZol reagent (ThermoFisher) according to manufacturer’s instructions. 3 ug of total cellular DNA is digested by BglII restriction enzyme, electrophoresed in an agarose gel, and digested DNA transferred to nylon membrane according to standard procedures. The probe for DNA hybridization was synthesized using DIG probe kit (Sigma-Aldrich) and used for overnight hybridization at 55C. Washing and processing of the blot was performed according to standard procedures.

### Chloroplast-enriched RNA fraction isolation, library construction, and sequencing

To isolate the chloroplast-enriched RNA fraction, we isolated chloroplasts from fresh leaves using the Minute^™^ Chloroplast Isolation Kit (Invent Biotechnologies, Plymouth, USA), following the manufacturer’s instructions. To isolate enough RNA, per sample, we isolated chloroplast from two preps of 250 mg of tobacco leaf. We isolated the chloroplast-enriched RNA fraction with the TRI Reagent (Sigma-Aldrich, St. Louis, USA) following the manufacturer’s instructions. To assess the purity of chloroplast-enriched RNA, between the whole-cell and chloroplast-enriched RNA fractions, we compared the abundance of two highly expressed miRNAs in tobacco leaves and observed that chloroplast RNA is enriched at 90% (Supplemental Table 2).

### Small library construction, sequencing and bioinformatic analysis

To collect the whole-cell RNA fraction, plantlets were harvested in three or five replicates and, after dissection, samples were immediately frozen in liquid nitrogen and kept at −80°C before RNA isolation. sRNA libraries were constructed using the RealSeq-AC miRNA Library Kit for Illumina sequencing (Somagenics, Santa Cruz, USA) using an input of 150 ng total RNA and 16 PCR amplification cycles. We size-selected sRNA libraries for the end product of ~150-nt using the SPRIselect Reagent (Beckman Coulter Life Sciences, Indianapolis, USA) magnetic beads. All libraries were quantified on a DeNovix apparatus (Wilmington, USA) using the Qubit dsDNA Assay Kit (Thermo Fisher Scientific, Waltham, USA) and we multiplexed libraries in 10 nM pools. Singleend sequencing was performed with 76 cycles (3 lanes). The sequencing was generated on an Illumina NextSeq 550 instrument (Illumina) at the University of Delaware DNA Sequencing and Genotyping Center.

We used cutadapt v3.4 (Martin, 2011) to preprocess sRNA-seq reads, removing the 3’ adapter and discarding trimmed reads shorter than 15-nt or longer to 40-nt. We mapped cleaned reads to *PDS1* (NCBI ID: XM_016610712.1) and *PDS2* (NCBI ID: XM_016642615.1) transcripts using ShortStack v3.8.5 (Johnson et al., 2016) with following parameters: -mismatches 0, -mmap u, -dicermin 15, -dicermax 40, and - mincov 1.0 reads per million (RPM). We used the ShortStack analysis report to identify a phasiRNA-generating feature over reads mapping each *PDS* gene for each study.

To visualize sRNA mapping to *PDS* genes, we converted mapping files into Bed Graph and Bed files using functions genomecov and bamtobed from bedtools v2.30.0 (Quinlan and Hall, 2010), respectively. We used the R package Sushi v1.24.0 (Phanstiel et al., 2014) to represent the position of transgenes over *PDS* genes and to visualize the coverage and read distribution of sRNAs over *PDS1* and *PDS2* genes.

To investigate properties of sRNA mapping to *PDS* genes, we investigated the read length distribution and the distribution of reads over *PDS* genes for constructs with a significant Phase Score. (i) We first summarized sRNA reads mapping to *PDS* genes into unique tags. We count the total number of reads per length and use the R ggpubr package (Wickham, 2016; Kassambara, 2020) to draw a bar plot of the read length distribution. For construction with a significant Phase Score, we had assigned the positions within the *PDS* genes to “phasing” bins to one of the 21 arbitrary bins, which repeat in 21-nt cycles (lines). We calculate the abundance of reads in each bin and visualize results on a radar plot. We followed the method described in the chapter “A Method to Discover Phased siRNA Loci” (Meyers and Green, 2009).

### Accession numbers

Sequence data of the genes used in this article can be found in Genbank under the following accession numbers: PDS1 (XM_016610712.1), PDS2 (XM_016642615.1).

## SUPPLEMENTAL DATA

*Supplemental Figure 1. Nuclear-encoded PDS genes organization, alignment and fragment insertion into the plastid genome*.

(A) Intron (blue lines) and exon (blue box) structure of the two nuclear-encoded *PDS* genes and cDNA fragment (red boxes) used for insertion into the plastid genome. (B) Alignment of *PDS1, PDS2* and the PTS transgene cDNA fragment. A 13 nucleotide sequence was inserted in the transgene fragment to discriminate plastid transgenes from the nuclear *PDS* genes. (C) A Southern blot showing the integration of the *PDS1* fragment and *aadA* transgenes at the expected location in the plastid genome. BglII restriction enzyme digest results in a ~5.8 kb fragment in the transplastomic lines while wild-type plants without transgene insertion carry an ~3.5 kb band. Note lack of the ~3.5 kb band in transplastomic lines indicating they are homoplasmic for transgenes insertion.

*Supplemental Figure 2. Mapping and abundance of PDS siRNAs in the PTS39-3 antisense RNA line*.

Low abundance siRNAs accumulate on both sense and antisense strands in the PTS39-3 line. Although abundance is low, the predominant read length is 21 nt (insert) and a moderate phasing score is observed for these siRNAs. The relative location of the transgene fragment (grey) in the *PDS* gene (light blue) are represented above the panels. The y-axis indicates the distribution of read abundance normalized in reads per million (RPM).

*Supplemental Figure 3. siRNAs in the PTS38 and PTS40 lines do not map to the PDS2 nuclear gene polymorphic regions*.

The figure shows the location of nucleotide polymorphisms (orange bars) between *PDS1* (left) and *PDS2* (right) gene regions. SiRNA reads distributed to each gene are shown for PTS38 (middle) and PTS40 (bottom) lines. Note the absence of siRNAs mapping to the 7 polymorphic positions of the *PDS2* gene. The y-axis indicates the distribution of read abundance normalized in reads per million (RPM).

*Supplemental Figure 4. Mapping of transgene derived small RNAs in purified chloroplast fractions of PTS38 and PTS40 lines*.

Mapping and accumulation of *PDS1* small RNAs (red, sense strand; blue, antisense strand) in the chloroplast fractions of PTS38 (A) and PTS40 (B) transplastomic lines. The relative location of the *PDS* transgene fragment (grey) in the nuclear-encoded *PDS1* gene (light blue) is represented above the panels. Note that the mapping pattern of small RNA reads from the PTS40 transgene differs from the whole-cell RNA fraction because sRNAs accumulate only on the positive strand of the transgene, and neither line accumulates any phasiRNAs in the chloroplast fraction. The y-axis indicates the distribution of read abundance normalized in read per millions (RPM).

*Supplemental Figure 5. shows that plastid-expressed dsRNA is processed to 21-nt phasiRNAs*.

(**A-B**) Mapping and accumulation of siRNAs in the MT91 and MT95 lines expressing the *Frankliniella occidentalis SN7F* gene from either convergent promoters (**A**) or a hairpin/loop RNA (**B**) transgene. Inserts show the length distribution of reads showing predominantly 21 nt siRNAs. The relative location of the transgene fragment in the *Snf7* gene is represented above the panels. The y-axis indicates the distribution of read abundance normalized in reads per million (RPM).

*Supplementary Table 1. Oligonucleotide probes used for qRT-PCR and northern blot experiments*.

*Supplementary Table 2. Purity of isolated chloroplast fractions as estimated from abundance of contaminating nuclear-encoded miRNAs*.

