## Supplementary Figure 1 for "Plastid double-strand RNA transgenes trigger small RNA-based gene silencing of nuclear-encoded genes"

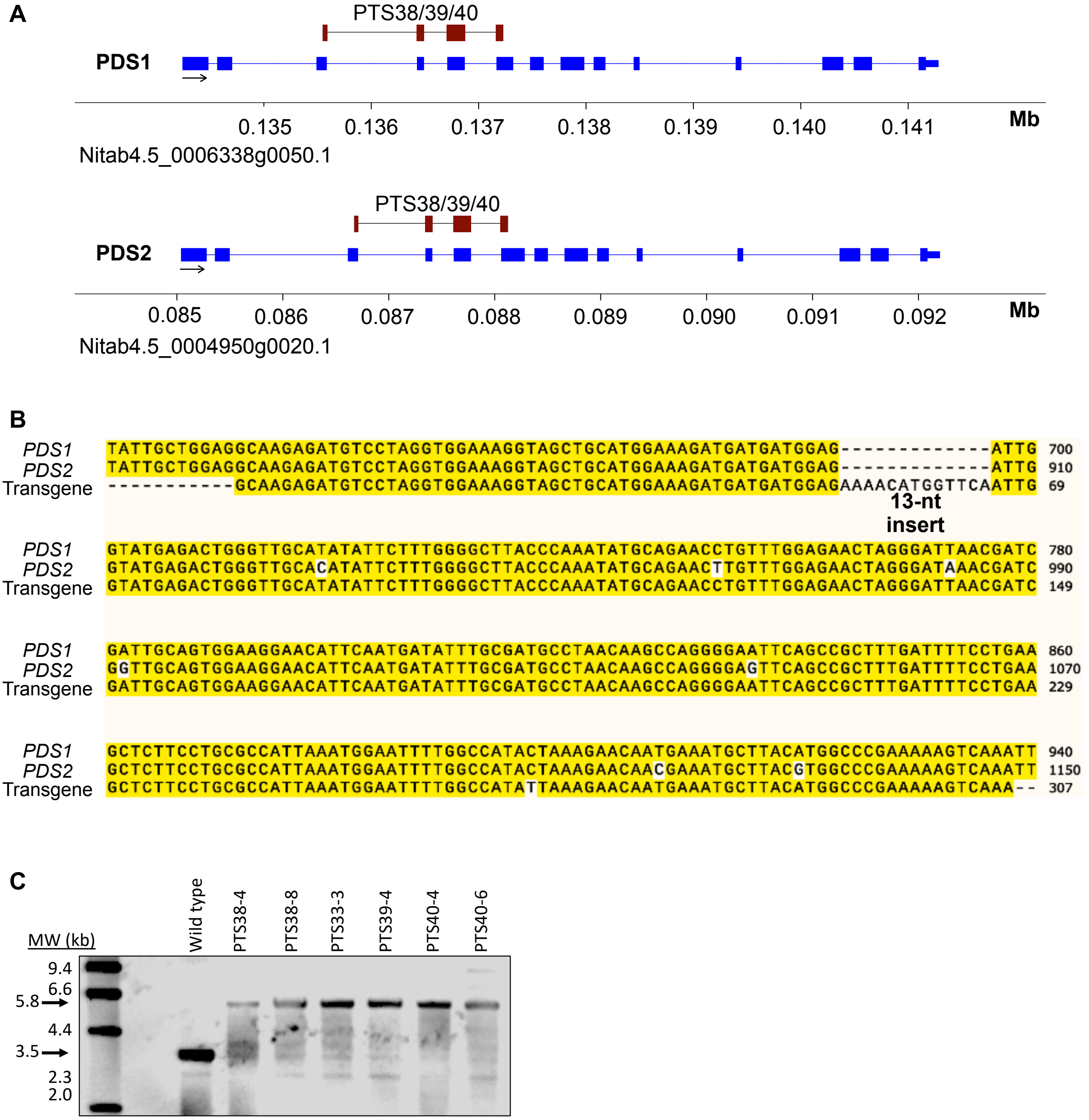


*Supplementary Figure 1. Nuclear-encoded PDS genes organization, alignment and fragment insertion into the plastid genome.*

(A) Intron (blue lines) and exon (blue box) structure of the two nuclear-encoded *PDS* genes and cDNA fragment (red boxes) used for insertion into the plastid genome. (B) Alignment of *PDS1*, *PDS2* and the PTS transgene cDNA fragment. A 13 nucleotide sequence was inserted in the transgene fragment to discriminate plastid transgenes from the nuclear *PDS* genes. (C) A Southern blot showing the integration of the *PDS1* fragment and *aadA* transgenes at the expected location in the plastid genome. BglII restriction enzyme digest results in a ~5.8 kb fragment in the transplastomic lines while wild-type plants without transgene insertion carry an ~3.5 kb band. Note lack of the ~3.5 kb band in transplastomic lines indicating they are homoplasmic for transgenes insertion.
