## Supplementary Figure 2 for "Plastid double-strand RNA transgenes trigger small RNA-based gene silencing of nuclear-encoded genes"

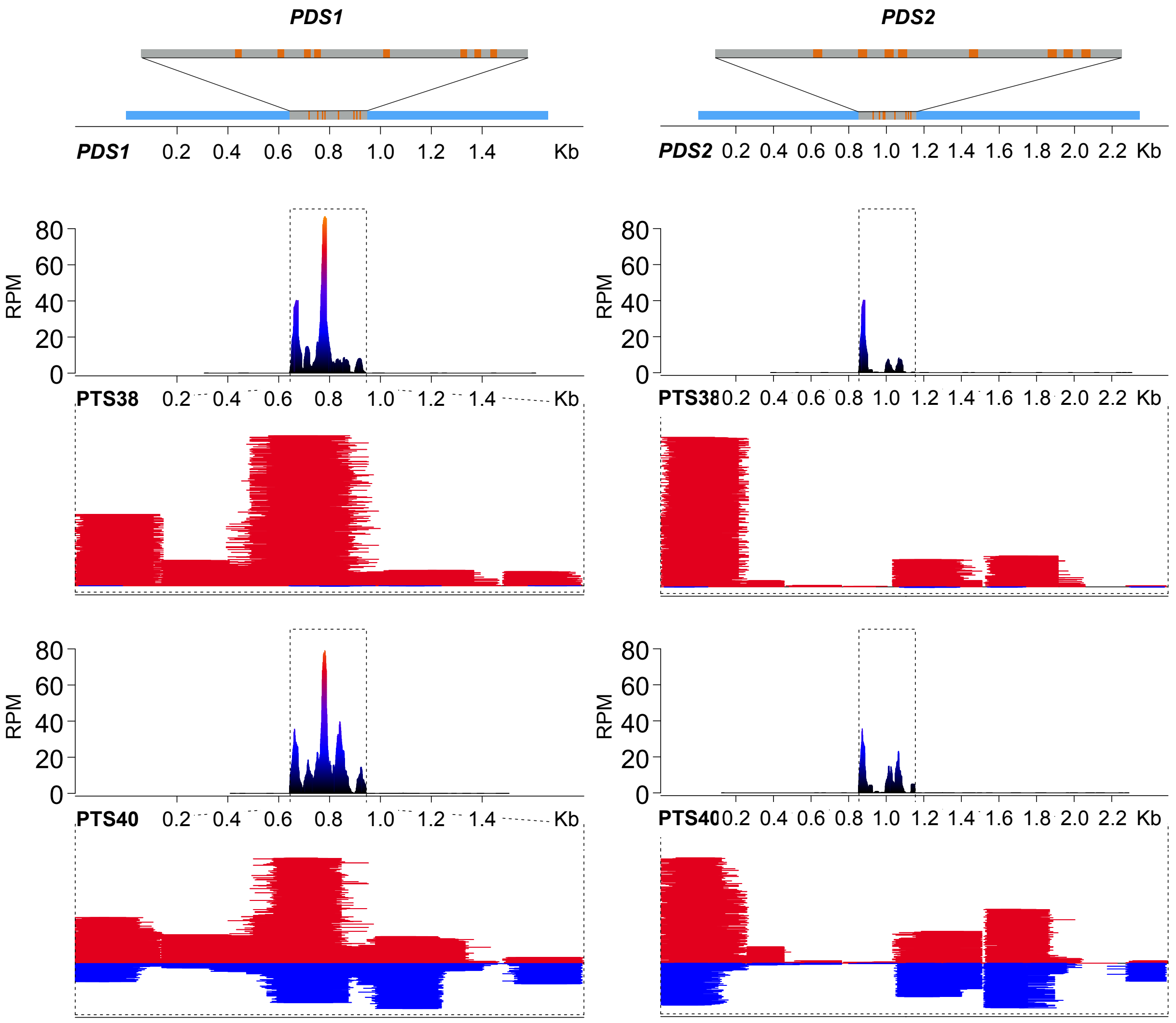


*Supplementary Figure 2. Mapping and abundance of PDS siRNAs in the PTS39-3 antisense RNA line.*

Low abundance siRNAs accumulate on both sense and antisense strands in the PTS39-3 line. Although abundance is low, the predominant read length is 21 nt (insert) and a moderate phasing score is observed for these siRNAs. The relative location of the transgene fragment (grey) in the *PDS* gene (light blue) are represented above the panels. The y-axis indicates the distribution of read abundance normalized in reads per million (RPM).
