## Supplementary Figure 3 for "Plastid double-strand RNA transgenes trigger small RNA-based gene silencing of nuclear-encoded genes"

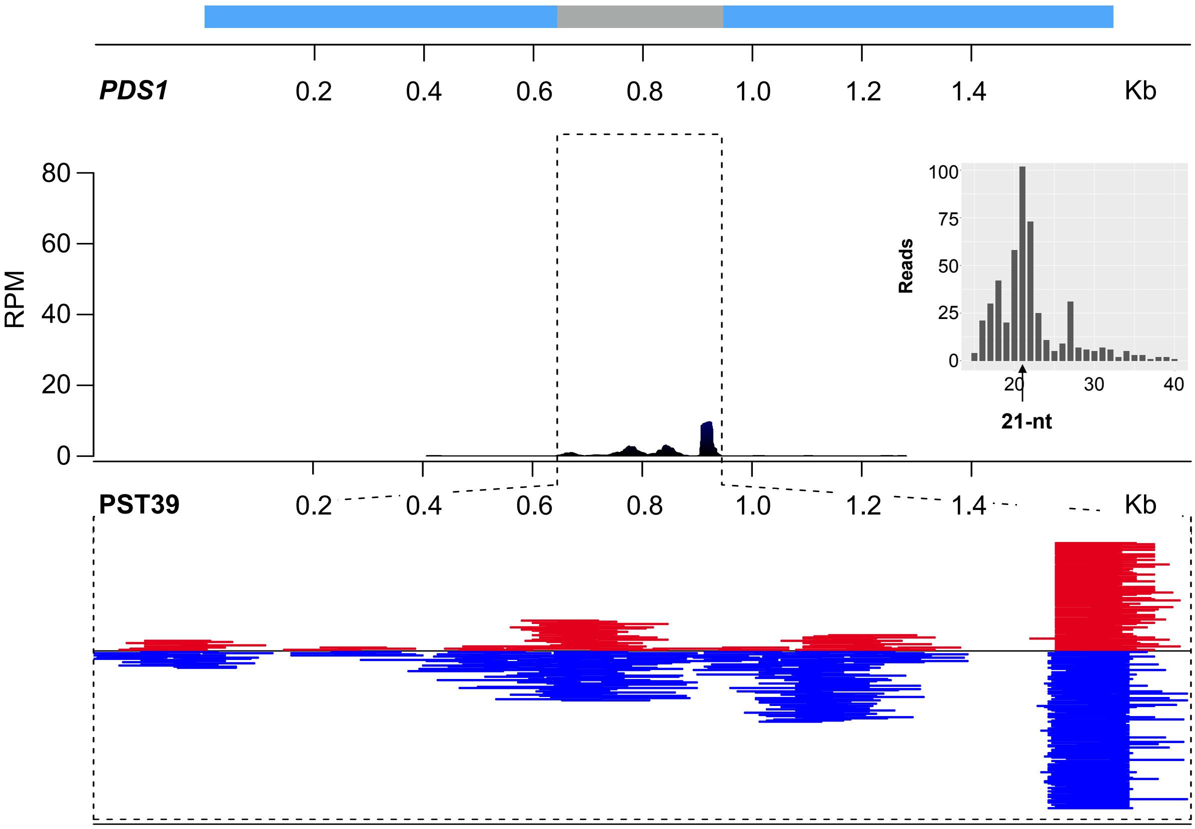


*Supplementary Figure 3. siRNAs in the PTS38 and PTS40 lines do not map to the PDS2 nuclear gene polymorphic regions.*

The figure shows the location of nucleotide polymorphisms (orange bars) between *PDS1* (left) and *PDS2* (right) gene regions. SiRNA reads distributed to each gene are shown for PTS38 (middle) and PTS40 (bottom) lines. Note the absence of siRNAs mapping to the 7 polymorphic positions of the *PDS2* gene. The y-axis indicates the distribution of read abundance normalized in reads per million (RPM).
