## Supplementary Figure 4 for "Plastid double-strand RNA transgenes trigger small RNA-based gene silencing of nuclear-encoded genes"

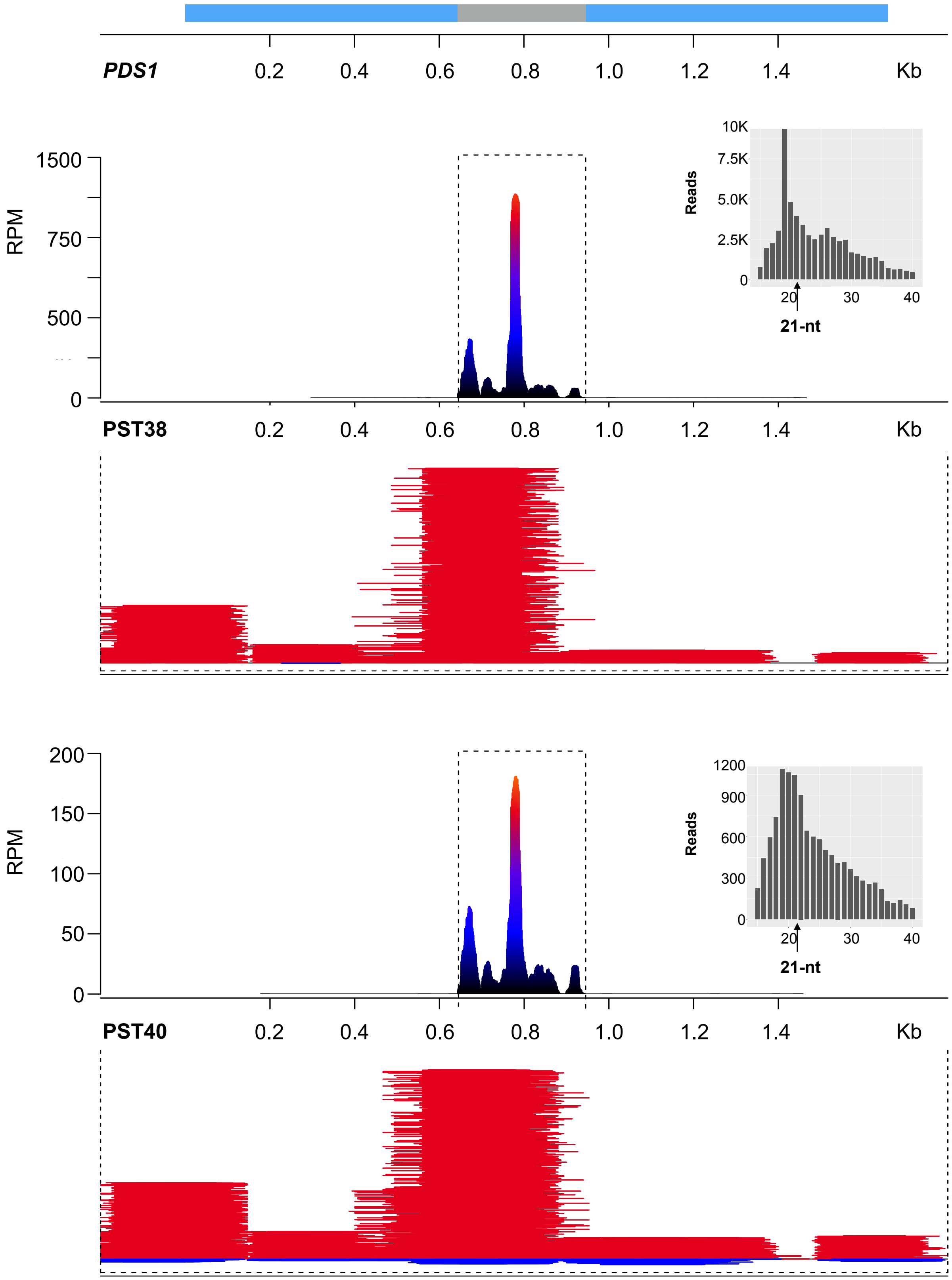


*Supplementary Figure 4. Mapping of transgene derived small RNAs in purified chloroplast fractions of PTS38 and PTS40 lines.*

Mapping and accumulation of *PDS1* small RNAs (red, sense strand; blue, antisense strand) in the chloroplast fractions of PTS38 (A) and PTS40 (B) transplastomic lines. The relative location of the *PDS* transgene fragment (grey) in the nuclear-encoded *PDS1* gene (light blue) is represented above the panels. Note that the mapping pattern of small RNA reads from the PTS40 transgene differs from the whole-cell RNA fraction because sRNAs accumulate only on the positive strand of the transgene, and neither line accumulates any phasiRNAs in the chloroplast fraction. The y-axis indicates the distribution of read abundance normalized in read per millions (RPM).
