## Supplementary Figure 5 for "Plastid double-strand RNA transgenes trigger small RNA-based gene silencing of nuclear-encoded genes"

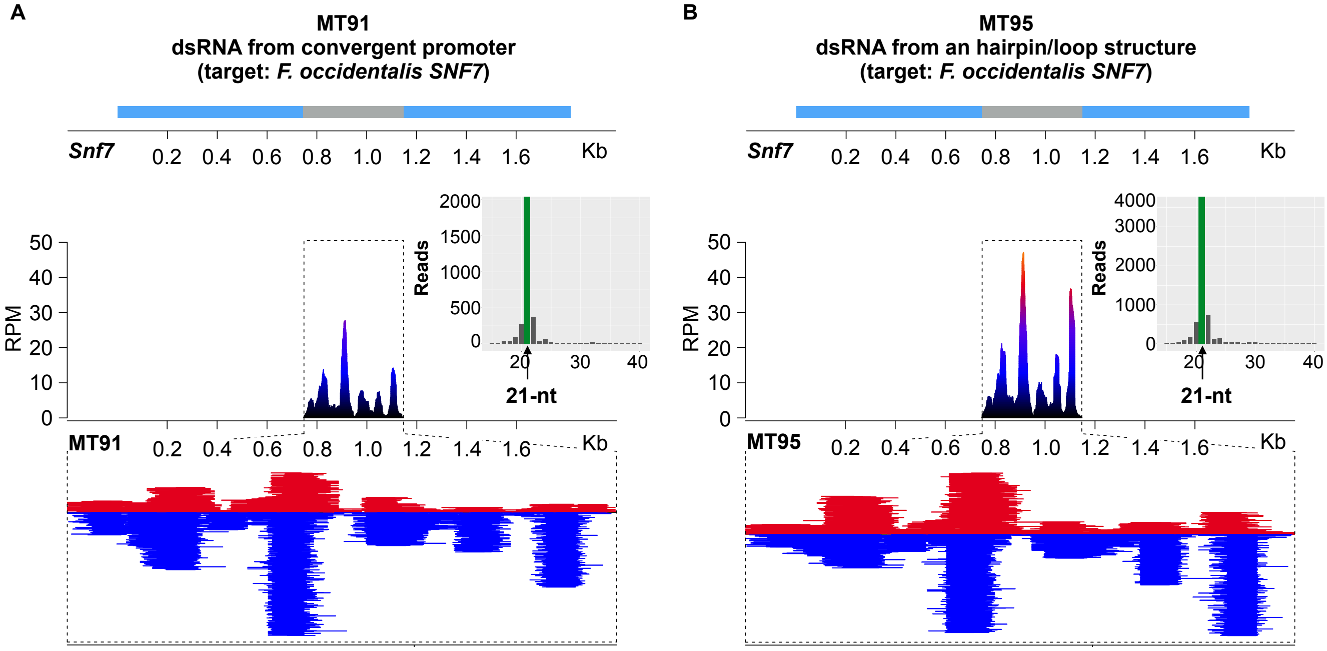


*Supplementary Figure 5. shows that plastid-expressed dsRNA is processed to 21-nt phasiRNAs.*
(A-B) Mapping and accumulation of siRNAs in the MT91 and MT95 lines expressing the Frankliniella occidentalis SN7F gene from either convergent promoters (A) or a hairpin/loop RNA (B) transgene. Inserts show the length distribution of reads showing predominantly 21 nt siRNAs. The relative location of the transgene fragment in the Snf7 gene is represented above the panels. The y-axis indicates the distribution of read abundance normalized in reads per million (RPM).
